# Molecular Imaging of the TGF-β Activating Integrin αvβ6 Detects Chronic Lung Allograft Dysfunction

**DOI:** 10.64898/2026.01.07.698265

**Authors:** Marlene Cano, Chandra S. Bathula, Fuyi Liao, Cristian Wieczorek Villas Boas, Yan Tao, Zhiyi Liu, Victoria Davis, Ricky Ebenezer, Allison Cannady, Dequan Zhou, Shunning Liang, Derek Byers, Alexander S. Krupnick, Daniel Kreisel, Zhiyu Dai, Buck Rogers, Andrew E. Gelman

## Abstract

TGF-β-activating integrins promote solid-organ fibrosis, suggesting their use as a molecular marker of disease. Chronic lung allograft dysfunction (CLAD), a progressive fibrotic complication that limits lung transplant survival, is driven by intragraft TGF-β activation. However, the expression patterns of TGF-β-activating integrins remain undefined in lung transplants. Single-cell RNA sequencing in a mouse CLAD model revealed high levels of the TGF-β-activating integrin αvβ6, which was mainly localized to fibrosis-associated Krt8^+^ transitional alveolar cells (AT1/2), while tolerant transplants lacked both αvβ6 expression and Krt8^+^AT1/2 cells. Molecular imaging with a newly developed positron emission tomography radiotracer specific for αvβ6, [^64^Cu]Cu-DOTA-A20-K16R, showed significantly higher uptake in CLAD versus tolerant transplants. In contrast, [^64^Cu]Cu-DOTA-A20-K16R allograft uptake was reduced by treatments that lowered αvβ6 expression and CLAD severity. Finally, [^64^Cu]Cu-DOTA-A20-K16R autoradiographic analysis on human explanted lungs with CLAD showed elevated activity that correlated with αvβ6 expression. Collectively, these findings demonstrate the potential utility of αvβ6 molecular imaging to detect CLAD pathogenesis.

## Introduction

Lung transplantation is the only life-saving option for end-stage pulmonary disease. Currently, the median survival of a lung recipient post-transplant is about six years, making it the worst life expectancy for all commonly transplanted solid organs. A major obstacle to long-term lung recipient survival is the development of chronic lung allograft dysfunction (CLAD), a fibrotic disease that has no effective treatments (1). The most common CLAD phenotype is Bronchiolitis Obliterans Syndrome (BOS), characterized by scarring of the small airways. The other phenotype, restrictive allograft syndrome (RAS), is identified by intra-alveolar fibrosis and elastosis. However, CLAD can also present with a varying mix of BOS and RAS pathological features (2).

Chronic TGF-β-activation has a well-established role in promoting tissue fibrosis. TGF-β, irrespective of isoform, is initially produced in an ‘inactive’ or latent form that is maintained by the non-covalent association of latency-associated peptide(s) (LAP).

Mechanical, oxidative, or proteolytic-mediated removal of LAP by TGF-β-activating molecules frees TGF-β to interact with cognate TGF-β receptors to induce profibrotic gene expression. For example, the heterodimeric TGF-β-activating integrins αVβ3, αVβ5, and αVβ6 recognize an arginine-lysine-aspartic acid (RGD) motif in the LAP, which, in turn, induces mechanical force that releases active TGF-β (3–6). In particular, αvβ6 plays a key role in activating TGF-β1 in solid organs. In the lung, αvβ6 is upregulated in patients with idiopathic pulmonary fibrosis (IPF). Integrin β6 (*Itb6*) gene ablation or αvβ6 pharmacological blockade inhibits radiation and bleomycin-induced pulmonary fibrosis (7, 8). However, whether αvβ6 is expressed during CLAD pathogenesis remains unclear.

A significant impediment to the development of effective CLAD therapies is the lack of methods for sensitive, early detection. CLAD is defined as the sustained loss of 20% or more of forced expiratory volume in 1 second (FEV1), a point at which the recipient is likely to have irreversible graft damage and has begun a path of continual functional decline. CLAD can also be diagnosed by transbronchial biopsy (Tbbx).

However, Tbbx suffers from low sensitivity due to the dispersed and patchy nature of CLAD fibrotic lesions, which can lead to false-negative diagnoses. Moreover, obtaining Tbbx requires bronchoscopy, which inherently carries risks of sedation, pulmonary bleeding, or pneumothorax in patients with already declining lung function. Finally, lung recipients can experience prolonged 10-20% FEV1 losses, but the lack of reliable diagnostic criteria makes it hard to confirm that this is a sign of impending disease.

Therefore, developing methods to detect CLAD development non-invasively could enable early diagnosis and guide early therapeutic interventions to prevent significant lung decline and potentially increase lung transplant graft survival.

Work by our group and others has led to the development and optimization of a positron emission tomography (PET) radiotracer ([^64^Cu]Cu-DOTA-A20-K16R) that detects αvβ6 expression on human and mouse cells (9). The probe’s specificity is derived from the A20 peptide of foot-and-mouth disease virus, which binds αvβ6 but not other TGF-β-activating integrins (10–12). Using a combination of single-cell RNA (scRNA) sequencing and proteomic analysis, we report elevated αvβ6 expression on Krt8^+^ transitional phenotype alveolar epithelial cells, which predominantly accumulate in mouse lung transplants with CLAD. PET-based [^64^Cu]Cu-DOTA-A20-K16R molecular imaging clearly distinguishes between CLAD and tolerant allografts and detects amelioratory responses to experimental CLAD treatments. Importantly, we also find that probing human explanted CLAD specimens with [^64^Cu]Cu-DOTA-A20-K16R results in elevated autoradiographic activity relative to control lung tissue.

## Results

### αvβ6^+^ transitional alveolar epithelial cells develop in allografts with CLAD

To analyze patterns of TGF-β-activating integrin expression in CLAD, we utilized an established mouse orthotopic lung transplant model (13–15) that leverages reported clinical observations of club cell dysfunction and injury in CLAD (16, 17) (Fig. 1).

**Figure 1.**
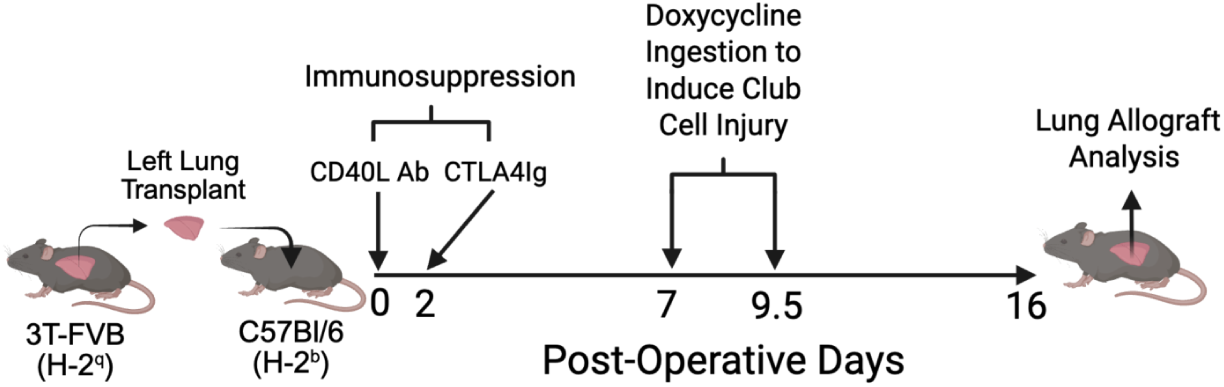
Induction of Club cell injury induces CLAD in mouse lung recipients. Left lungs from 3T-FVB mice are transplanted into C57Bl/6 recipient mice and treated with CD40L Ab and CTLA4Ig on POD 0 and POD 2, respectively, to induce allograft tolerance. Doxycycline ingestion from post-operative day (POD) 7 to POD 9.5 induces club cell injury that triggers CLAD by POD 16. Recipients that do not ingest Doxycycline remain tolerant to their allografts.

Immunosuppressed C57Bl/6 (H-2^b^) mice received major-histocompatability-complex mismatched left lungs encoding three transgenes (3T-FVB, H-2^q^); a reverse tetracycline activator gene driven by the club cell secretory protein promoter, a Cre recombinase gene under the control of the reverse tetracycline activator, and a lox-P-activated diphtheria toxin A gene (14). Doxycycline ingestion induces transgene-mediated club cell injury, leading to loss of allograft tolerance, characterized by a mixture of airway and alveolar epithelial injuries that reflect a mixture of BOS and RAS phenotypes found in humans (13–15). In contrast, if transplant recipients are not induced to undergo club cell injury, lung allografts remain tolerant (14).

Single-cell RNA sequencing (scRNA) analysis of tolerant and CLAD allografts produced uniform manifold approximation and projection (UMAP) plots, which facilitated the annotation of 31 cell phenotypes within clusters of stromal, endothelial, epithelial, and immune cells (Sup. Fig. 1). Examination of the CLAD allograft epithelial cell cluster revealed a high proportion of alveolar epithelial cells with a mixed phenotype (AT1/2) that expressed transcripts characteristic of alveolar type 2 cells (AT2), such as Lamp3, Sftpb, and Sftpc, as well as alveolar type 1 cells (AT1), including Ager, Cav1, and Hopx (18, 19) (Figs. 2A-D). CLAD AT1/2 cells also displayed elevated levels of Cldn4 and Krt8 transcripts. Notably, the accumulation of Krt8^+^ alveolar epithelial transitional cells has been reported in models of lung fibrosis and in humans with idiopathic lung fibrosis (20–22), consistent with alveolar epithelial injury observed in human RAS (2) and our model (15). Consistent with prior work showing that AT2 cells rapidly proliferate and differentiate into AT1 cells to repair alveolar epithelium (23, 24), proliferating AT2 cells (Prolife_AT2) were much more prevalent in CLAD allografts, indicating an ongoing injurious response. Finally, epithelial cell trajectory analysis inferred that CLAD AT1/2 cells are transitional intermediates between AT2 cells and AT1 cells (Fig. 2E).

**Figure 2.**
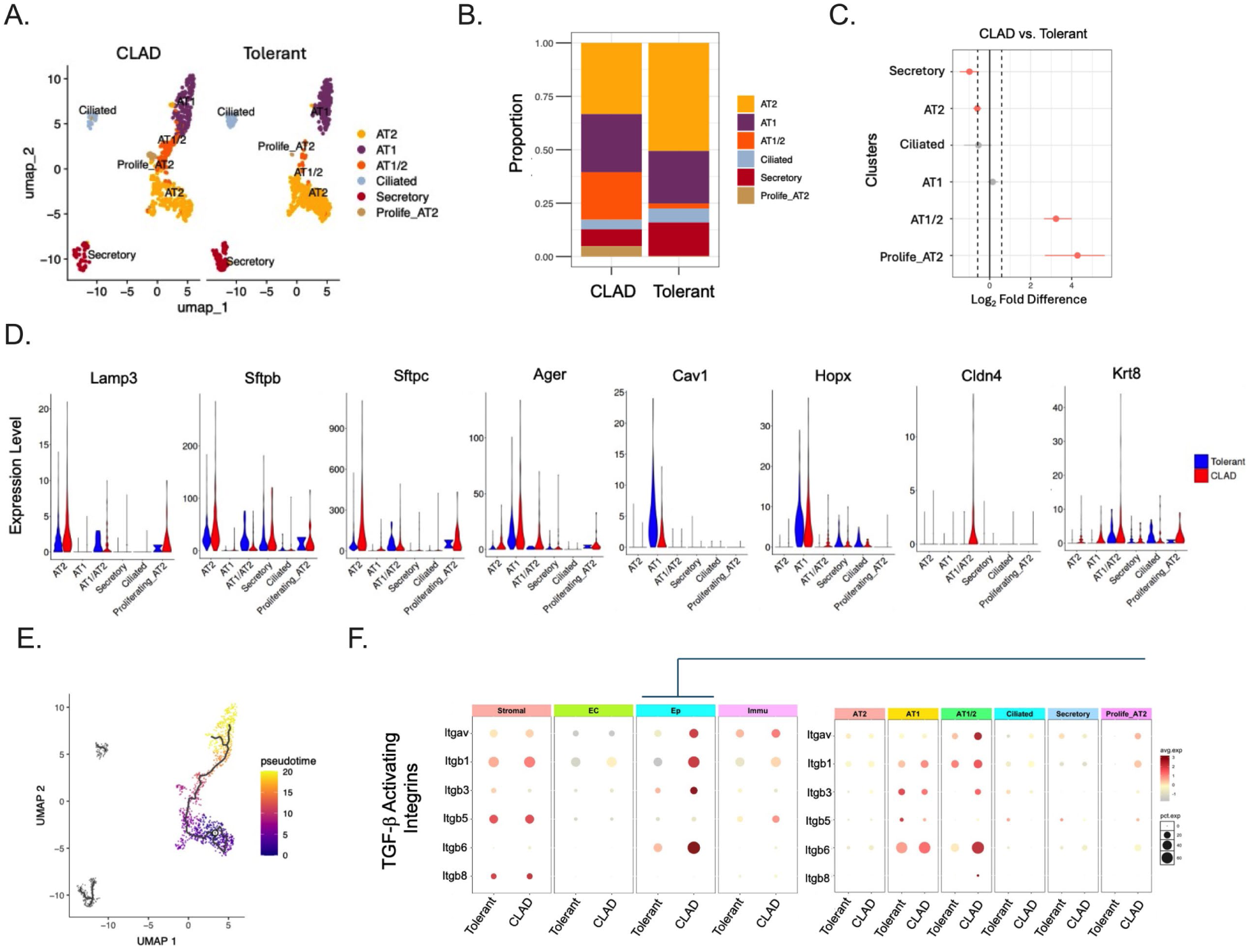
αvβ6 transcripts are enriched in transitional alveolar epithelial cells. POD 16 CLAD and tolerant allograft tissue analyzed by single-cell RNA analysis. (A) Uniform Manifold Approximation and Projection (UMAP) on 21,778 cells derived from 3 tolerant and 3 CLAD allografts, with epithelial clusters shown and analyzed for the (B) relative proportionality and (C) Log2 fold differences for each cell phenotype shown with confidence intervals. (D) Violin plots shown for canonical genes that mark AT1 (Lamp3, Sftpb, Sftpc), AT2 (Ager, Cav1, Hopx), and AT1/2 (Cldn4, Krt8) cells. (E) Epithelial cellular trajectory analysis conducted on CLAD allografts. (F, left panel) TGF-β activating integrin dot blot expression analysis for all lung cells (F, right panel) with a breakdown on epithelial cell phenotypes. Prolife_AT2; Proliferating AT2 cells.

Regarding the well-established role of TGF-β-activating integrins in promoting fibrosis (25), we assessed transcript levels of integrin Itgav (αv) subunit family members in CLAD and tolerant allografts (Fig. 2F). While αv transcripts were detected in stromal and immune cell compartments irrespective of allograft rejection, epithelial cells from allografts with CLAD had the highest levels. Additionally, CLAD epithelial cells exhibited co-accumulation of transcripts encoding αv heterodimeric subunit partners Itgb1 (β1), Itgb3 (β3), and Itgb6 (β6). Differentiating for specific epithelial cell phenotypes, AT1/2 cells produced the highest relative amounts of transcripts encoding for TGF-β-activating integrins, with Itgb6 levels being the most pronounced, indicating that transitional alveolar epithelial cells are the primary source of αvβ6 expression in CLAD lungs. To further explore this observation, we performed flow cytometric analysis on cell suspensions isolated from CLAD and tolerant allografts. αvβ6^+^ cells showed evidence of an AT2 origin, as gauged by high EpCAM (CD326) and partial surfactant protein C (Sftpc) expression, and were approximately six times more numerous in CLAD allografts relative to allografts from tolerant recipients (Figs. 3A-C). Consistent with these findings, we observed greater αvβ6 expression in CLAD allografts by immunohistochemistry (Fig. 3D).

**Figure 3.**
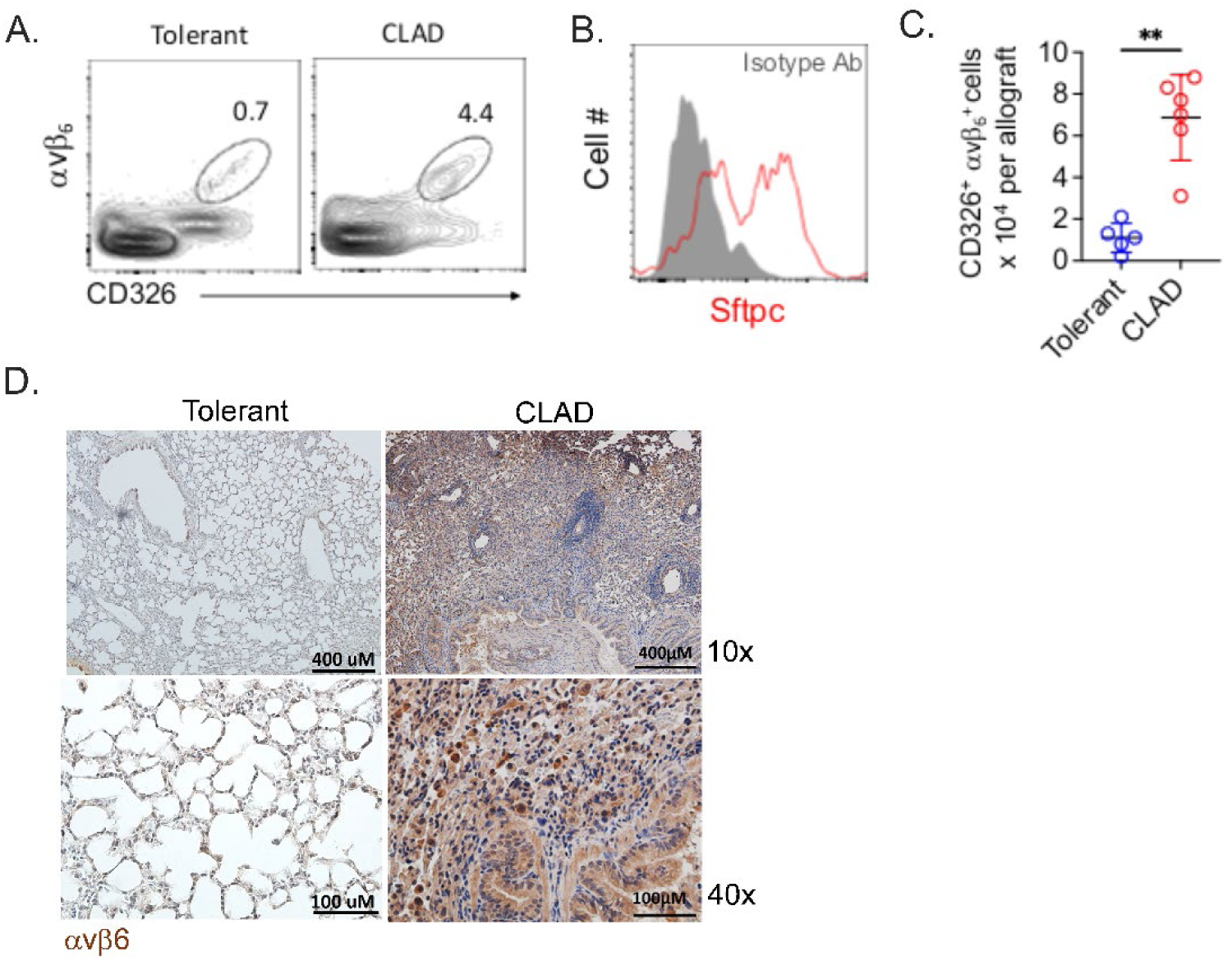
Elevated avβ6 protein expression in lung allografts with CLAD. (A) Flow cytometric analysis of POD 16-tolerant and CLAD allografts with (A) representative contour plots gated on live CD45-allograft cells (N=5/group) and (B) Intracellular Sftpc expression analysis on CLAD allograft CD326+ αvβ6+ cells. The histogram shown is representative of three independent experiments. (C) Plot of POD 16 allograft numbers of live CD45-CD326+ αvβ6+ cells shown with a mean ± standard deviation from the mean for N=5 per group. Representative immunohistochemical stains with αvβ6 specific antibodies with indicated magnification and scales (N≥4/group).

### Intragraft molecular imaging of αvβ6 detects CLAD development

In mouse lung recipients with tolerant allografts and allografts that developed CLAD, we performed positron emission tomography (PET) imaging with the αvβ6-specific radiotracer [^64^Cu]Cu-DOTA-A20-K16R (9). On post-operative day (POD) 16, lung recipients were injected intravenously with 11.1 MBq (300 μCi) of [^64^Cu]Cu-DOTA-A20-K16R and imaged by computerized tomography (CT), followed by static PET scans for 1 hour. Compared with tolerant lung allografts, allografts with CLAD showed increased radiotracer uptake (Fig. 4A). Intragraft [^64^Cu]Cu-DOTA-A20-K16R activity measured by mean standard uptake value (SUV Mean) revealed that CLAD allografts had nearly 3-fold more mean activity relative to tolerant lung allografts (Fig. 4B). To assess probe target specificity, a 100-fold excess of A20 peptide (cold probe) was administered just before [^64^Cu]Cu-DOTA-A20-K16R radiotracer imaging of lung recipients with CLAD. Cold-probe challenge abolished [^64^Cu]Cu-DOTA-A20-K16R uptake in CLAD lungs (Fig. 4A-B). We next performed hematoxylin and eosin (H&E) and Masson’s trichrome staining on imaged lung allografts (Fig. 4C). In contrast to tolerant lung transplants, allografts with high [^64^Cu]Cu-DOTA-A20-K16R activity or that received cold probe exhibited histological evidence of CLAD, as noted by the appearance of obliterative lesions and pleuroparenchymal fibrosis. Collectively, these data indicate that αvβ6 molecular imaging distinguishes allografts with CLAD from tolerant allografts.

**Figure 4.**
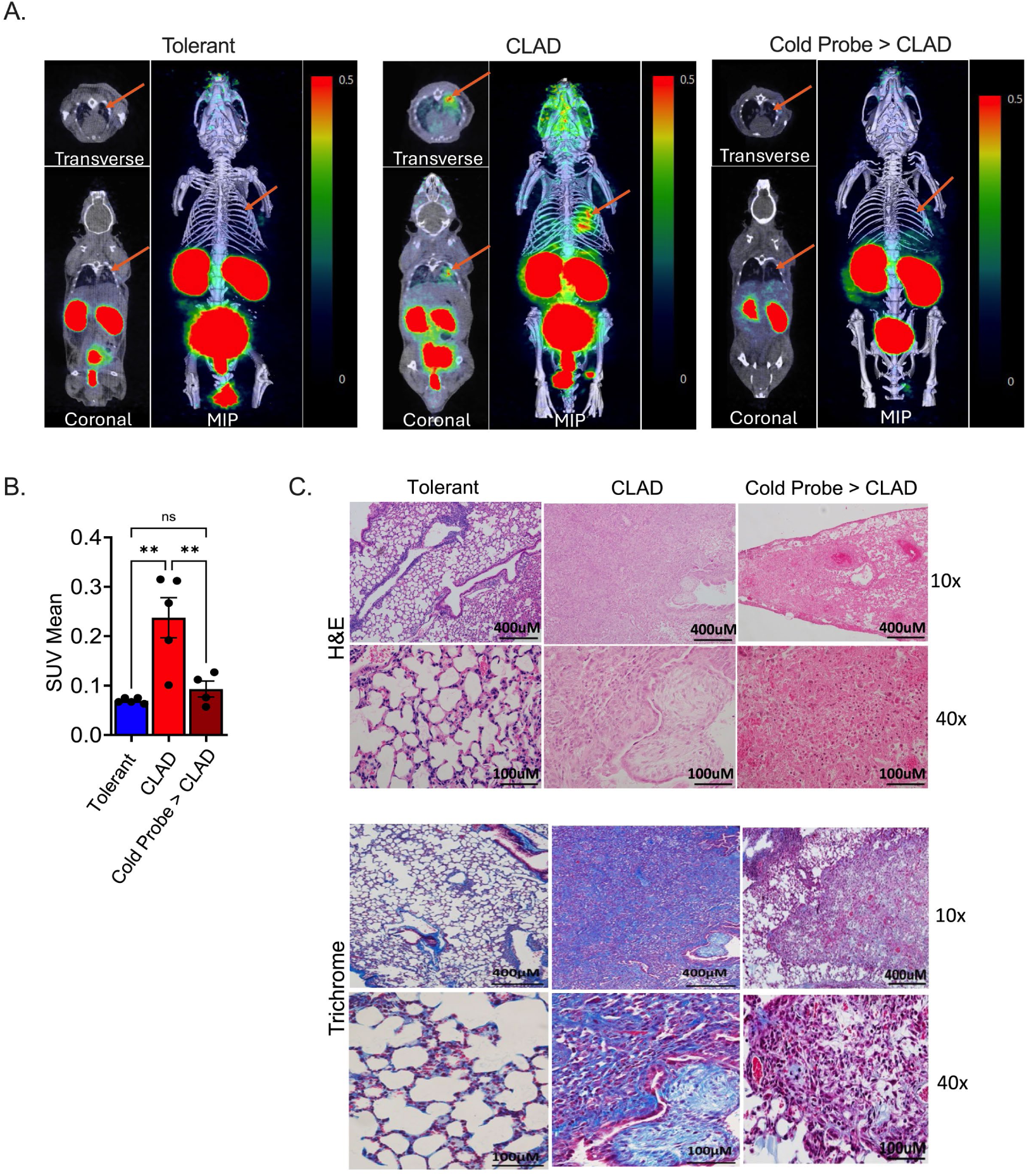
PET/CT imaging with αvβ6-specific radiotracer reveals high uptake in CLAD allografts. (A, left and middle panels) [^64^Cu]Cu-DOTA-A20-K16R PET/CT scans of POD 16 recipients with lungs that are tolerant (N=5) or have CLAD (N=5). (A, right panel) lung recipients with CLAD that received cold peptide blockade (Cold Probe > CLAD) on POD 16 prior to PET/CT imaging (N=4). Images shown are representative scans for each group shown in transverse, coronal and maximum intensity projections (MIP). (B) Plots of radiotracer intragraft uptake for each transplant assessed by SUV Mean shown with group mean ± standard error of the mean (SEM). A two-sided Mann-Whitney U-test was conducted to evaluate significance, where **p<0.01. Each data point represents a single transplant. (C) POD 16 allograft H&E and Trichrome staining (N≥4/group). Scale bars: 400 μm for 10x, 100 μm for 40x.

### Inhibition of CLAD severity reduces αvβ6 expression

Recent work by our group has demonstrated that treating lung recipients with CCL2-neutralizing antibodies (Ab) to reduce profibrogenic CCR2^+^ monocyte graft infiltration, or administering CD8α-specific Abs to deplete CD8^+^ T cells, inhibits CLAD pathogenesis (14, 15). Regarding these previous observations, we asked whether either of these treatments would inhibit [^64^Cu]Cu-DOTA-A20-K16R allograft uptake. CCL2 Ab neutralization or CD8^+^ T cell depletion inhibited [^64^Cu]Cu-DOTA-A20-K16R allograft activity compared to respective control Ig administration (Figs. 5A, B). Consistent with our previous work, both treatments markedly attenuated histological signs of CLAD severity (Fig. 5C). Importantly, αvβ6 expression was also attenuated, irrespective of the CLAD severity-reducing treatment (Figs. 6A, B). Together, these findings suggest that αvβ6 molecular imaging can monitor the effectiveness of anti-CLAD therapies.

**Figure 5.**
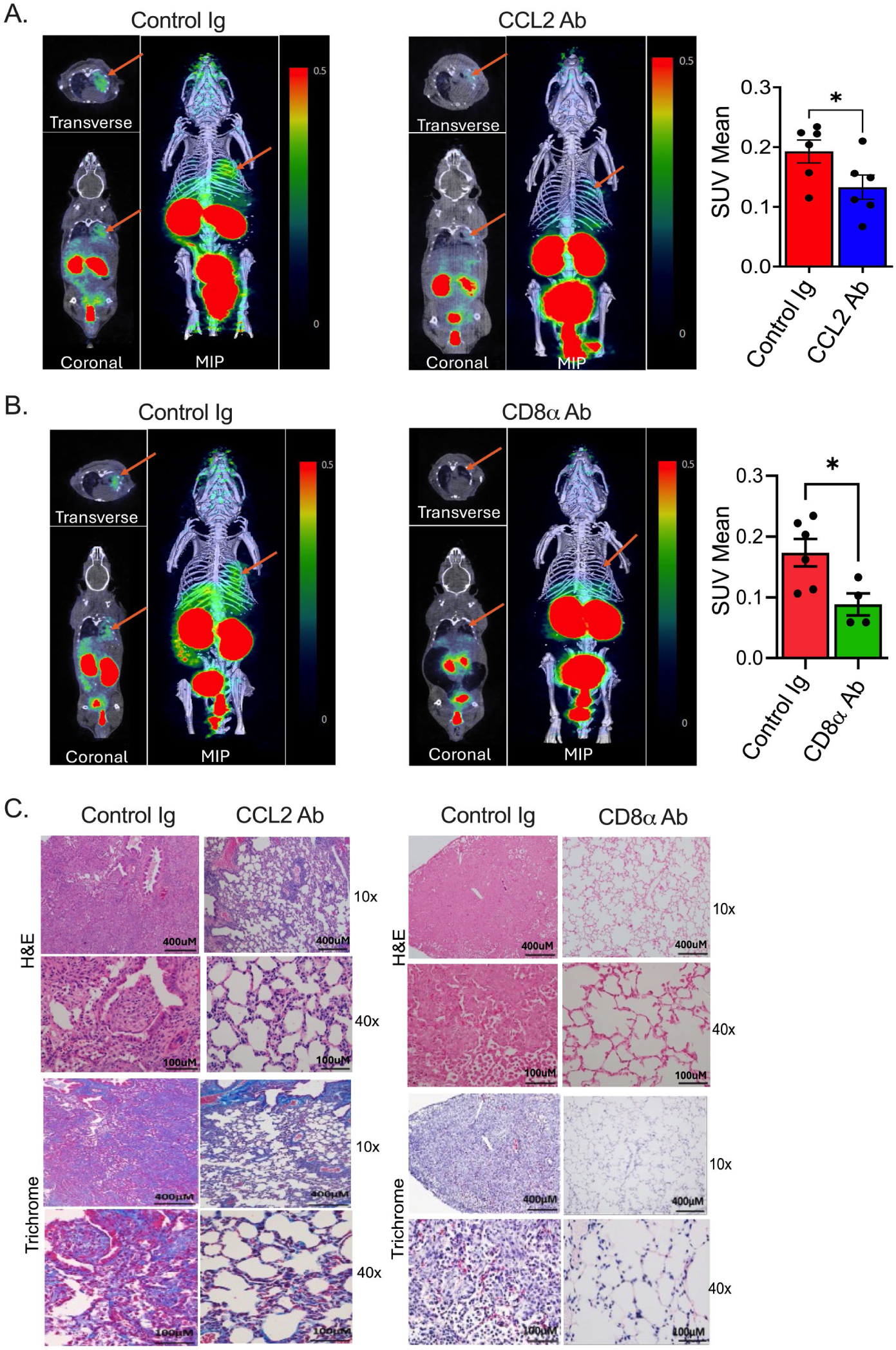
Inhibiting CLAD severity reduces [^64^Cu]Cu-DOTA-A20-K16R allograft uptake. Lung recipients that underwent club cell injury and received either CCL2 neutralizing Abs, CD8α depleting Abs, or respective Control Ig injections and then underwent [^64^Cu]Cu-DOTA-A20-K16R PET/CT imaging on POD 16. Scans shown (A, B) are representative transverse, coronal and maximum intensity projections (MIP) images for each indicated treatment with N≥4 for each group. The accompanying plots show SUV Mean for each transplant with group mean ± SEM. A two-sided Mann-Whitney U-test was conducted to evaluate significance, where *p<0.05. (C) POD 16 representative H&E and trichrome stains for indicated treatments (N≥4/group) with scale bars: 400 μm for 10x, 100 μm for 40x.

**Figure 6.**
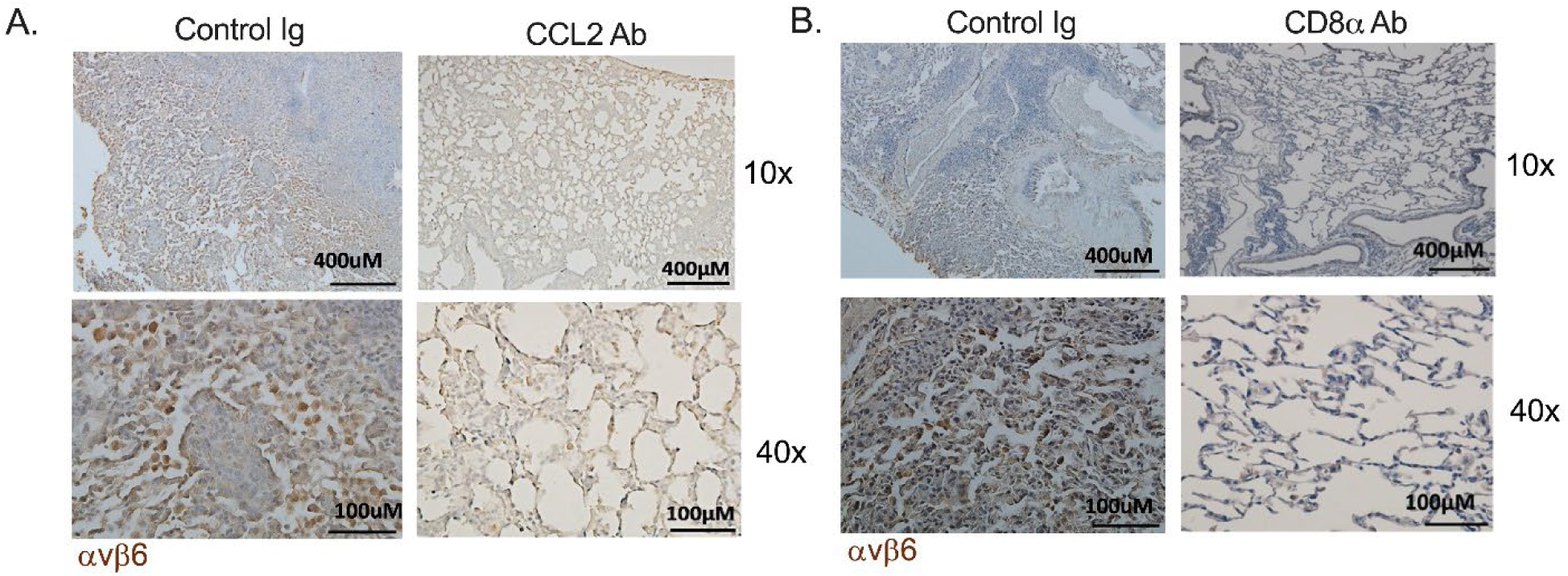
Reducing CLAD severity with CCL2 or CD8α Ab treatment inhibits αvβ6 expression. αvβ6 immunohistological staining of allografts from recipients that underwent club cell injury and received (A) CCL2 neutralizing Abs, (B) CD8α depleting Abs, respectively, and (A, B) Control Ig injections followed by POD 16 [^64^Cu]Cu-DOTA-A20-K16R PET/CT imaging. Images shown are representative of N≥4 per group. Scale bars: 400 μm for 10x, 100 μm for 40x.

### Explanted human lung transplants with CLAD express αvβ6

To assess whether αvβ6 molecular imaging could be useful for detecting CLAD in human lung recipients, we performed [^64^Cu]Cu-DOTA-A20-K16R autoradiography on control lung tissue and explanted lung transplants from recipients diagnosed with CLAD (Figs. 7A, B). Compared to control lung tissue obtained from donor allografts prior to transplantation, CLAD showed higher levels of radiotracer binding, which could be significantly reduced by pre-incubation with an excess of cold probe.

**Figure 7.**
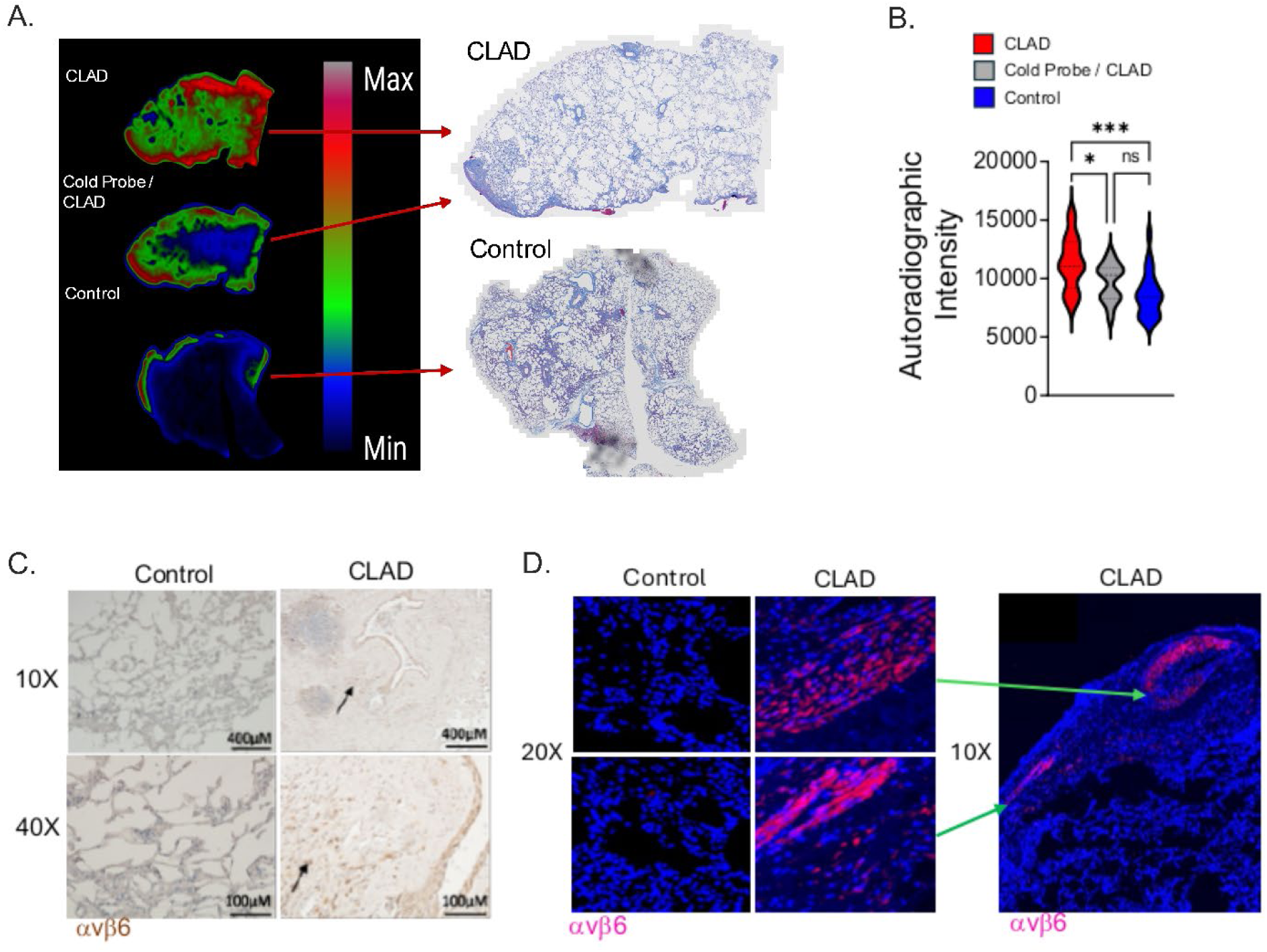
Human lung transplants with CLAD bind the [^64^Cu]Cu-DOTA-A20-K16R radiotracer. (A, Left Panel) Representative autoradiographic images following incubation of [^64^Cu]Cu-DOTA-A20-K16R with human explanted lung tissue sections from a patient with CLAD and Control lung tissue from a donor lung obtained prior to transplantation. Preincubation with an excess of a cold probe sharply reduces autoradiographic activity (Cold Probe/CLAD). (A, Right Panel) corresponding trichrome-stained lung sections that were incubated with radiotracer. Images are representative of independently conducted experiments, with (B) violin plots showing combined data from 3 explanted CLAD transplants with or without cold-probe blockade and 3 control lungs per group. Two-sided Mann-Whitney U t-test *p<0.05, ***p<0.001. Representative αvβ6 (C) immunohistochemical and (D) immunofluorescent staining of human explanted lungs with CLAD and control lungs (N=3/group). Arrows denote clusters of cells expressing αvβ6. Scale bars: 400 μm for 10x, 100 μm for 40x.

Immunohistochemical and immunofluorescent staining revealed numerous αvβ6-expressing cells scattered within the allograft parenchyma and close to the pleural membrane (Figs. 7C, D). In contrast, control lung tissue exhibited minimal αvβ6 expression.

## Discussion

We report that elevated [^64^Cu]Cu-DOTA-A20-K16R radiotracer activity is indicative of CLAD pathogenesis in lung transplants from mice and humans. It is important to note that other probe designs have been studied for non-invasive imaging of αvβ6 expression in cancer and pulmonary fibrosis models. A considerable number of these probes were based on a cysteine-knot peptide (26–28), a monomeric or trimeric cyclic nonapeptide called trivehexin (29, 30). While trivehexin peptides exhibit high specificity for αvβ6, they have limited ability to chelate several important radionuclides, including ^64^Cu, thereby reducing their utility for PET imaging (29). The radiotracer used in this study is based on the A20 peptide, originally isolated from a phage-display library, which identified the DLXXL sequence as critical for αvβ6 specificity (11, 12, 31). This A20 peptide was later modified by bi-terminal pegylation and a lysine-to-arginine substitution at position 16 (K16R) to increase probe αvβ6-dependent cell-binding affinity and reduce non-target solid organ uptake (9, 32). In a mouse model of focal radiation-induced lung fibrosis, in which αvβ6 expression also serves as a biomarker of disease development, we recently demonstrated that [^64^Cu]Cu-DOTA-A20-K16R uptake was restricted to irradiated regions (9).

Consistent with clinical observations, we previously reported that TGF-β activation drives CLAD pathogenesis in the 3T-FVB lung transplant model (15, 33, 34). Notably, TGF-β activation is increased in CLAD allografts but not in the circulating blood of lung recipients with CLAD or in tolerant allografts (15). Our scRNA transcript analysis suggested that high levels of αv integrins on the alveolar epithelium of CLAD allografts could be potent activators of TGF-β in the allografts. While Itgav and Itgb6 transcripts were sharply upregulated in epithelial cells, there was also co-enrichment of Itgb1 and Itgb3, indicating that αvβ1 and αvβ3 contribute to intra-graft TGF-β activation. Like αvβ6 inhibition, antagonism of αvβ1 has been shown to reduce disease severity in bleomycin-induced lung fibrosis in mice (35). The role of αvβ3 in lung fibrogenesis is less clear, although it has been shown to promote fibroblast contraction in mice (25). The impact of TGF-β-activating integrin expression in human lung fibrotic diseases is currently being evaluated in drug trials. In a recent phase 2 clinical trial with IPF patients, treatment with the dual selective inhibitor of αvβ6 and αvβ1, PLN-74808, not only improved lung function but also reduced collagen deposition (36), a direct target of TGF-β-mediated signaling (37). In a phase 2a clinical trial evaluating treatment with the αvβ6-neutralizing antibody BG00011 in IPF patients, there was a significant reduction in TGF-β-mediated signaling in BAL cells (38). However, in a subsequent phase 2b trial, the study was terminated early due to severe adverse events, which may have been related to the need to retain some αvβ6 activity to maintain pulmonary homeostasis (39). While these observations indicate a more complex role for αvβ6, they do not rule out its potential as a biomarker for human pulmonary fibrotic disease. In support of this, we found that elevated αvβ6 expression correlated with high [^64^Cu]Cu-DOTA-A20-K16R autoradiographic activity in explanted human lungs with CLAD, whereas control lungs lacked both αvβ6 expression and autoradiographic activity.

ScRNA epithelial cluster profiling indicated that αvβ6 expression was almost exclusively limited to transitional AT1/2 cells in CLAD allografts. Interestingly, CLAD AT1/2 cells exhibited increased levels of Keratin 8 (Krt8) transcripts, an intermediate filament protein that helps maintain cytoskeletal mechanical strength and regulates cell differentiation. Krt8 is also an established marker of transitional alveolar epithelial cell phenotypes observed in mouse lung fibrosis models and in patients with IPF (20).

Elegant studies by Wang and colleagues have shown that Krt8 is essential for the generation of transitional alveolar epithelial cells in mice (40, 41). Remarkably, mouse transitional alveolar epithelial cells share features with Krt8^+^ basaloid transitional alveolar cells found in the lungs of IPF patients, as they appear arrested in a non-determinative, highly senescent state and lack a robust capacity to differentiate into AT1 cells (20, 21). Previous research has also demonstrated that Itgb6 transcripts are upregulated in Krt8^+^ transitional alveolar epithelial cells in lungs with IPF and that inhibiting early TGF-β signaling reduces Krt8 expression in cultured alveolar epithelial cells (21). In line with these observations, we previously demonstrated that intragraft TGF-β activation is low in tolerant allografts (15), possibly reflecting the lack of AT1/2 cell accumulation in this setting. This suggests that Krt8 might be essential for αvβ6 expression. To this end, overexpressing Krt8 in cell lines has been shown to increase Itgav transcript levels (42). However, to date, no reports have documented a requirement for Krt8 in the upregulation of Itgb6. Lastly, scRNA studies in human lung transplants have not clearly demonstrated enrichment of Krt8^+^ transitional alveolar epithelial cells during CLAD pathogenesis, despite reporting elevated transcript levels of genes with known profibrogenic activity in alveolar epithelial cell clusters (43, 44). One possible reason for the lack of detection is the relative rarity of a fulminant RAS-CLAD phenotype compared to BOS. Since RAS primarily involves alveolar-centric injury, transitional alveolar epithelial cells are expected to be significantly more abundant in RAS specimens than in BOS specimens. Additionally, unintended selection bias for airway fibrotic lesions could make it more difficult to detect such cells, highlighting the need to phenotype CLAD in future scRNA analyses.

To further investigate the potential of αvβ6 molecular imaging, we performed [^64^Cu]Cu-DOTA-A20-K16R CT/PET imaging following treatments that we previously demonstrated reduce CLAD severity (14, 15). Initially, CCR2-neutralizing antibodies were administered to prevent the development of CCR2^+^ monocyte-derived alveolar macrophages (Mo-AM) (15). We observed a significant decrease in [^64^Cu]Cu-DOTA-A20-K16R intragraft activity, which was associated with less histological evidence of CLAD and decreased αvβ6 allograft expression. Regarding this treatment, our group and others have shown that Mo-AMs, which accumulate in response to lung epithelial injury, promote CLAD and bleomycin-induced lung fibrosis in mice (15, 45). After lung transplantation in humans, donor-derived alveolar macrophages are replaced by Mo-AM, a process that seems to be accelerated by primary graft dysfunction, a form of ischemia-reperfusion injury (46). Importantly, primary graft dysfunction is a risk factor for CLAD (47), indicating that future therapies targeting Mo-AM development could be monitored using [^64^Cu]Cu-DOTA-A20-K16R PET-based imaging. In our second set of studies, we used CD8α Abs to deplete CD8^+^ T cells in lung recipients undergoing CLAD pathogenesis and observed reduced [^64^Cu]Cu-DOTA-A20-K16R intragraft activity and allograft αvβ6 expression. While the reasons for the reduction in αvβ6 expression following either of these treatments are unknown, we have observed that Mo-AM expand intragraft resident-memory IFN-γ^+^ CD8^+^ T cells, which, in turn, are required to promote CLAD by causing epithelial injury (15). Interestingly, in a recent study of influenza virus-induced lung fibrotic injury in mice, Mo-AM and lung resident IFN-γ^+^ CD8^+^ T cells were observed clustering around Krt8^+^ transitional alveolar epithelial cells (48). Conversely, depletion of pulmonary-resident CD8^+^ T cells reduced the levels of Krt8^+^ transitional epithelial cells and prevented alveolar epithelial damage. Anti-thymocyte immunoglobulin (ATG) depleting therapies have been used in human lung transplant recipients to slow CLAD progression (48). Although multicenter, randomized controlled trials are needed to identify predictors of favorable responses to ATG treatment, several studies suggest that early intervention may improve outcomes, highlighting the need for facile, non-invasive methods to detect early-onset CLAD development.

In summary, we identified patterns of TGF-β-activating integrin expression in lung transplants and found that high αvβ6 expression is limited to Krt8^+^ transitional alveolar epithelial cells that accumulate in CLAD allografts. We then used this observation to evaluate the potential of αvβ6 molecular imaging for detecting CLAD. However, our study has some limitations. First, it remains unclear whether targeting αvβ6 function alone would be sufficient to prevent CLAD. While we observed a correlation among αvβ6 expression, [^64^Cu]Cu-DOTA-A20-K16R uptake, and CLAD development, future studies should explore the use of TGF-β-activated integrin inhibitors in combination with radiotracer imaging. In this context, a recent clinical trial involving eight IPF patients demonstrated that an αvβ6 cystine-knot peptide radiotracer competed with the PLN-74809 αvβ6/αvβ1 dual-specific inhibitor for αvβ6 receptor occupancy (49).

Nevertheless, additional work will be needed to dissect the patterns of αvβ6 expression in human lung recipients and to further validate this approach for detecting CLAD development.

## Methods

### Sex as a biological variable

The orthotopic lung transplant model used male or female donor lungs transplanted into male recipients. Male recipients were utilized in this study due to the use of the penile vein for injections of immunosuppressants. However, sex was not considered a biological variable in this study.

### Radiochemical synthesis of [^64^Cu]Cu-DOTA-A20-K16R

The peptide 1,4,7,10-tetraazacyclododecane-1,4,7,10-tetraacetic acid (DOTA)-PEG28-NAVPNLRGDLQVLAQRVART-PEG28, here abbreviated as DOTA-A20-K16R, was synthesized and characterized by AnaSpec (Fremont, CA, USA). The DOTA-A20-K16R was dissolved in HPLC-grade water at 1 nmol/μL, fractionated in 10 or 20 μL, and stored at -20°C until use. Whatman 60 Å silica gel thin-layer chromatography (TLC) plates and a Bioscan 200 imaging scanner (Bioscan, Inc., Washington, DC, USA) were used for quality control. All solvents and reagents were purchased from Sigma-Aldrich (St. Louis, MO, USA) or Fisher Scientific (Pittsburgh, PA, USA) and used as received unless stated otherwise. HPLC-grade water was used to prepare all solutions and buffers. The ^64^Cu was produced from ^64^Ni(p,n)^64^Cu nuclear reaction on enriched ^64^Ni on a TR-19 biomedical cyclotron (Advanced Cyclotron Systems, Inc., Canada) at Washington University in St Louis, followed by an automated system using standard procedures for purification (50, 51). A stock solution of ^64^Cu in 0.1M HCl was diluted ten-fold with 0.1 M Ammonium Acetate (NH4OAc) pH 5.5. An aliquot containing 148 MBq (4 mCi) of ^64^Cu was transferred to a microtube and diluted to 92 μL with 0.1 M Ammonium Acetate (NH4OAc) pH 5.5, followed by addition of 8 μL of DOTA-A20-K16R (8 nmol) such that the final reaction volume was 100 mL. The reaction mixture was incubated at 95°C for 15 minutes (min). The radiochemical purity of the ^64^Cu-labeled peptide was evaluated by radio-TLC with a mobile phase of 50 mM Diethylenetriamine pentaacetate (DTPA) pH 6.0. The [^64^Cu]Cu-DOTA-A20-K16R remained at the origin, while free [^64^Cu]CuCl2 moved with the solvent front.

### Small animal PET/CT imaging

Mice were injected intravenously with approximately 11.1 MBq (300 μCi) of [^64^Cu]Cu-DOTA-A20-K16R (diluted in 0.9% sterile saline solution, total volume of 100 μL). For blocking experiments, cold K16R peptide (250 µg in 100 µL sterile saline) was injected intravenously 10 min prior to radiotracer administration. The mice were imaged by CT followed by static PET scans at one hour after radiotracer administration on a nanoScan PET/CT scanner (Mediso Medical Imaging Systems, Budapest, Hungary). Static images were collected for 45 min and reconstructed using the Order Subset Expectation Maximization Method with the Mediso software (Mediso Medical Imaging Systems).

Region of interest (ROI) was selected based on co-registered anatomical CT images and the radioactivity associated in those areas. Quantification (SUVmean) was measured using the Imalytics PreClinical software 3.1.1.4 (Gremse-IT, Aachen, Germany).

### Autoradiography

Paraffin-embedded human lung tissue sections were deparaffinized in three consecutive xylene changes (5 min each) and rehydrated through graded ethanol to distilled water. Antigen retrieval was performed by heating the slides in a steamer for 20 min in preheated citrate buffer (pH 6.0; Sigma-Aldrich, C9999). The slides were allowed to return to room temperature, and the sections were washed in 1x wash buffer (Dako, S3006). The slides were incubated in dual endogenous enzyme block (Dako S2003) for 10 min to quench the endogenous peroxidase activity. 1% bovine serum albumin in PBS containing 0.1% Tween-20 was used to block the nonspecific binding. The tissues were incubated with 11.1 MBq/mL of [^64^Cu]Cu-A20-K16R for 2 hours at room temperature and rinsed four times with purified water (MilliQ). The A20 peptide (NAVPNLRGDLQVLAQKVART) was used as a blocking agent. Samples were exposed to a phosphor plate overnight and revealed in a phosphor screen (GE Typhoon FLA 9500 Variable Mode Laser Scanner) at 50-micron resolution. The mean intensity of autoradiography for a tissue sample was obtained using ImageJ.

### Single-Cell RNA Sequencing

POD 16 lung allograft cells were loaded onto a 10 X Genomics Chromium Single Cell Controller to generate barcoded single cells for constructing single-cell cDNA libraries. ScRNA sequencing was performed on a HiSeq 4000. Cell Ranger Pipeline v 9.0.0 was used for demultiplexing and counting. The 10X mouse transcriptome mm10-2020-A served as the reference genome. The R package Seurat (v 5.2.0) was employed for data preprocessing and visualization. Initially, cells with fewer than 200 genes or complexity (Log 10 (genes/UMI)) less than 0.0.8 were removed. Genes expressed in fewer than 5 cells were discarded. Doublets were identified using a two-layer approach as previously described: first, scDblFinder (v 1.16.0) was used for doublet discrimination, using 30 principal components for clustering with a resolution set at 0.5. Uniform Manifold Approximation and Projection (UMAP) was used for cluster visualization. Marker genes for each cluster were identified using the FindAllMarkers function. A small cluster of cells showing no canonical marker gene expression and a high mitochondrial gene ratio was considered low quality and removed. The remaining cells were reclustered, resulting in 31 distinct clusters. Cell identities were assigned based on previously published canonical marker genes with the AddModuleScore function in Seurat, according to LungMAP. Quality control steps to filter the count matrices included removing cells with over 5% mitochondrial RNA content, cells expressing more than 8,000 genes (due to the high risk of multiple cells in the barcoded GEM), and cells expressing fewer than 500 genes. Normalization and variance stabilization of raw counts were performed using SCTransform, and cell cycle scores were computed and regressed out together with percentage mitochondrial reads (52).

The normalized R object was then used for subsequent clustering and differential expression analysis. Briefly, clusters were annotated into major cell populations, each major cell type was subsetted, re-normalized, and subjected to PCA, UMAP embedding, clustering, and differential expression analysis. Monocle 3 was used to construct the epithelial cell trajectories.

### Mice and Lung Transplant CLAD Model

Animal studies were approved by the Institutional Animal Care and Use Committee at Washington University in Saint Louis. 3T mice on the FVB background have been previously described and were used as donors for all studies (14, 15). To induce allograft acceptance, recipients received intraperitoneal (i.p.) injections of CD40L Abs (250 μg; clone MR1, Bio-X-Cell) on POD 0 and human recombinant CTLA4 Ig (200 μg; Bio-X-Cell) on POD 2. Club cell injury was triggered by DOX ingestion via food (625 mg/kg chow; ENVIGO) and water (2 mg/mL, MilliporeSigma) for 2.5 days. CD8^+^ T cell-depleting Abs (500 μg, clone 53-6.7; Bio-X-Cell) were administered i.p. on POD 6 and 11. CCR2-neutralizing Abs (200 µg; clone C2-Mab 6, Bio-X-Cell) were injected intravenously once daily from POD 6-15. Rat IgG1 control isotype Abs (clone TNP6A7, Bio-X-Cell) were used for both studies.

### Hematoxylin & Eosin (H&E) and Trichrome Histological Staining

Paraffin-embedded slide sections were deparaffinized and rehydrated in alcohol and stained with Hematoxylin (Millipore Sigma, Cat# H9627) for 4 min and Eosin (Millipore Sigma, Cat# 318906) for 1 min before washing, dehydration, clearing, and mounting. Trichrome staining was performed using a Masson Trichrome Kit (Millipore Sigma cat# HT15) according to the manufacturer’s recommendations.

### Immunohistochemistry staining

For immunohistochemical analysis, paraffin sections were serially blocked at room temperature with BLOXALL Blocking solution (Vector Laboratories, SP-6000) 10 min, Blocking Fc (PK-6101, Vector Laboratories) 20 min, and Avidin/Biotin Blocking serum for 15 min each (SP-2001, Vector Laboratories). Sections were then stained with 1:200 αvβ6 antibodies (bs-5791R, Bioss for mouse tissue; AB00891-23.0, Absolute Antibody for human tissue) overnight at 4°C. For secondary antibodies, paraffin sections were serially incubated at room temperature with a drop of biotinylated antibody and Vectastain ABC Reagent (PK-6101, Vector Laboratories) each for 30 min, then with ImmPACT DAB Reagent (SK-4105, Vector Laboratories) for 1-2 min. Each step with 3 times PBS wash. Counterstaining and Mounting were applied before permanent mounting.

### Immunofluorescence staining

For immunohistochemical analysis, paraffin sections were blocked with 5% BSA at room temperature for 45 min. Sections were then stained with 1:200 αvβ6 antibodies (bs-5791R, Bioss for mouse tissue; AB00891-23.0, Absolute Antibody for human tissue) overnight at 4°C. For secondary Ab-mediated immunofluorescent visualization, we used 1:700 goat anti Rabbit cy5, 60 min at room temperature. Autofluorescence quenching kit (SP-800, Vector Laboratories) was applied 3 min at room temperature. Each step with 3 times PBS wash.

### Flow Cytometric Analysis

Lung tissue was digested and prepared for single-cell suspension as previously described (53). Live/dead cell staining was performed with the Zombie Violet Fixable Viability Kit (Biolegend). Cell surface staining was conducted with the following antibodies: CD45 (30-F11,BD), CD326 (clone G8.8, Thermofisher), and Integrin alpha V beta 6 (clone 53a.2, Thermofisher). Intracellullar staining was accomplished with Cytofix/Cytoperm kit (BD Biosciences) using polyclonal Rabbit anti-mouse/human STFPC ( Invitrogen, Cat# PA5-71680) or polyclonal Rabbit IgG Isotype Control (Thermofisher, Cat# 02-612) followed by staining with Goat anti-rabbit IgG Alex Flour 488 (Thermofisher, Cat# A78953).

### Statistical analysis

Statistical analysis was performed using GraphPad Prism (version 10.6). For comparisons between three groups, ANOVA was used with a Tukey’s HSD post-hoc test, and for non-pairwise comparisons, the Mann-Whitney U non-parametric Test was employed where indicated. A P-value < 0.05 was considered statistically significant.

### Study Approval and Human Lung Tissues

Animal experiments were conducted in accordance with an approved IACUC protocol (Washington University, 19-0827). Human participants provided written informed consent in accordance with the Washington University School of Medicine Institutional Review Board for Human Studies protocol (ID #201103213, #201012829). Explanted tissue specimens obtained from human lung recipients who were diagnosed with CLAD before retransplantation. Control lung tissue was healthy donor lung tissue obtained prior to transplantation.

