## Supplemental Figure for "Molecular Imaging of the TGF-β Activating Integrin αvβ6 Detects Chronic Lung Allograft Dysfunction"

Supplemental Figure 1

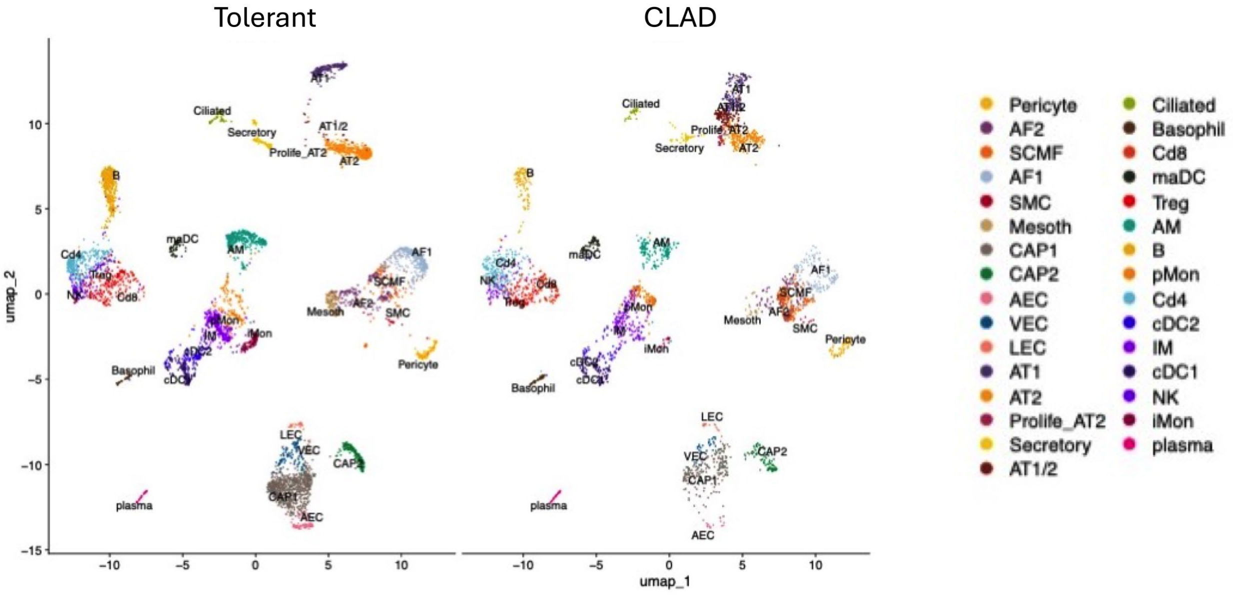


**Supplemental Figure 1. ScRNA analysis of tolerant and CLAD allografts reveals differences in alveolar epithelial cell clusters.** POD 16 CLAD and tolerant allograft tissue analyzed by single-cell RNA analysis. (A) Uniform Manifold Approximation and Projection (UMAP) of 21,778 cells derived from 3 tolerant and 3 CLAD allografts.
